# mV2G: a multiomic atlas for tissue-specific variant-to-gene prioritization

**DOI:** 10.64898/2026.08.11.743997

**Authors:** Dean Y. Zhang, Hufeng Zhou, Maya U. Sheth, Andreas R. Gschwind, Jesse M. Engreitz, Xihong Lin, Hongbo Liu

## Abstract

The multiomic Variant-to-Gene (mV2G, https://mv2g.hbliulab.org) is a comprehensive atlas that integrates diverse functional genomic evidence to prioritize tissue-specific variant-to-gene (V2G) associations. While genome-wide association studies (GWAS) have identified millions of associations between genetic variants and diseases, translating these findings into biological mechanisms remains challenging because >90% of variants reside in noncoding regions. Existing V2G resources provide complementary regulatory evidence but are fragmented and often lack tissue-specific interpretation. To address this challenge, we constructed the mV2G atlas by integrating 24 types of functional genomic evidence across 50 human tissues, including molecular quantitative trait loci, enhancer–gene predictions, three-dimensional chromatin interactions, and experimental validation. The atlas contains 188,634,118 evidence-supported V2G pairs involving 13,618,039 variants and 69,521 genes. We further developed a unified tissue-specific V2G prioritization framework and prioritized 1,530,420 high-confidence functional V2G pairs involving 1,131,316 unique variants, with 87% exhibiting tissue-specificity. The mV2G atlas provides searchable variant- and gene-centered interfaces, an interactive browser for visualizing variants, target genes, cis-regulatory elements, and chromatin states, as well as downloadable datasets. By integrating complementary regulatory evidence into a unified framework, mV2G provides an accessible resource for interpreting the functional and phenotypic impact of genomic variation in relevant tissues for human diseases.

**GRAPHICAL ABSTRACT:** 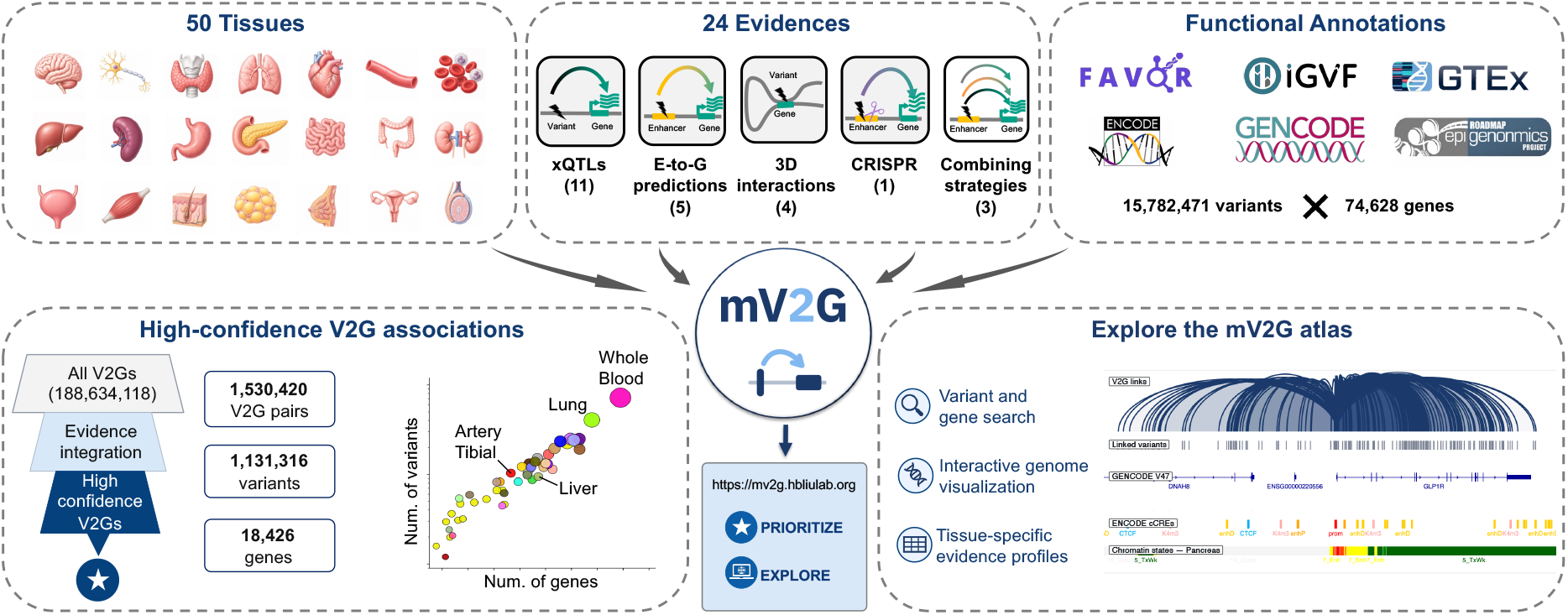

## INTRODUCTION

Genome-wide association studies (GWASs) have identified millions of associations between genetic variants and complex traits and diseases (1). However, translating these associations into mechanistic insights remains a major challenge because more than 90% of these disease-associated variants reside in non-coding regions, a well-known “variant-to-function” problem (2). In particular, these non-coding variants often regulate genes in a tissue-specific manner: a variant may influence gene expression in a particular tissue while having little or no regulatory effect in another (3-5). Determining which genes are regulated by these variants, and in which tissues those regulatory effects occur, remains one of the central challenges in human genetics (4,6). Thus, comprehensive collection and prioritization of variant-to-gene (V2G) relationships across human tissues are important for systematic exploration of the tissue-specific effects of non-coding variants on gene regulation and human disease processes.

Numerous experimental and computational approaches have been developed to infer V2G relationships, including molecular quantitative trait locus (xQTL) analyses (4,5,7-13), enhancer-gene linking methods (14-19), chromatin interaction assays (20-23), functional perturbation studies (16), and tissue-agnostic integrative frameworks (24-26). Each approach provides complementary evidence supporting V2G relationships. However, because these methods differ in the biological signals they measure, underlying assumptions, experimental designs, and tissue coverage, no single approach provides a complete picture of the regulatory landscape. As a result, V2G evidence is dispersed across numerous databases, web portals, publications, and supplementary datasets, hindering comprehensive evaluation and interpretation of causal variants and V2G relationships in human diseases.

Recent studies have developed several evidence combining strategies, including cS2G (24), Open Targets (25), and GeneHancer (26), to identify disease-causal V2G pairs. But these strategies generally lack a tissue-specific context, limiting their ability to identify tissue-specific functional V2G associations. Many of these resources also focus only on disease-associated loci, fine-mapped candidates, or coding variants, despite noncoding variants accounting for the vast majority of genetic variation (2). Furthermore, several published integrative frameworks lack dedicated database interfaces and evidence-centered exploration tools, limiting accessibility and direct evaluation. Our recent study in the human kidney demonstrated the promise of the tissue-level multiomic V2G (mV2G) strategy for prioritizing tissue-specific disease-causal variants and genes (27). However, a resource extending this strategy across diverse human tissues has remained unavailable.

To this end, we developed the mV2G atlas, a comprehensive tissue-specific V2G database that integrates diverse multiomic datasets to prioritize high-confidence regulatory variants and their target genes across 50 human tissues. The current release combines more than 20 types of tissue-level variant-to-gene associations, and together, these features enable systematic prioritization and exploration of tissue-specific V2G relationships across diverse biological contexts.

## MATERIAL AND METHODS

### Annotations of variants, genes, and variant-to-gene pairs

The genetic variants, annotated by the Functional Annotation of Variants - Online Resource (FAVOR) database, were derived from the Impact of Genomic Variation on Function (IGVF) Data Portal (curated sets ID: IGVFDS5361GWAK) (28-30). FAVOR provided an initial total of 1,052,126,302 globally unique variants. The common variants were retained by FAVOR’s “COMMON” category and a minor allele frequency of more than 0.01. Any variants with reference or alternative alleles more than 50 base pairs (bp) were filtered out. For multi-allelic variants, only the alternative allele with the highest frequency was kept. After filtering, a unique set of 15,782,471 variants remained for further analysis. The set of annotated genes was derived from Release 47 of Genome Coding Denotation (GENCODE) (31), with 74,628 unique genes, including 19,355 protein-coding genes and 55,273 non-protein-coding genes. Both variants and genes had Grch38/hg38 information. To identify potential V2G pairs, we created a 1 Mbp window around each variant and retained all genes that overlapped with that region, resulting in 425,576,568 potential V2G pairs as a reference. Variant rsIDs and Ensembl gene identifiers were used as standardized identifiers to ensure consistency across datasets and resource versions.

### V2G evidence collection across 50 human tissues

The V2G evidence collection was performed in 50 human tissue types that are widely used in population-based xQTL studies, such as the Genotype-Tissue Expression (GTEx) project (4). Totally, 24 types of evidence were collected and organized into five major categories, including four categories that capture tissue-specific evidence: xQTLs, enhancer–gene predictions, three-dimensional (3D) chromatin interactions, and CRISPR-based experimental validation; and the fifth category, tissue-agnostic V2G combining strategies, providing evidence applied across all tissues (**Table 1**). For xQTL datasets, we retained statistically significant V2G associations reported by the original studies, preserving their tissue specificity and matching them to our curated set list. For enhancer-gene prediction resources, variants overlapping a predicted enhancer were linked to the enhancer’s assigned target gene in their respective tissues. For 3D chromatin interaction datasets, variants within one interaction anchor were linked to genes whose transcription start site (TSS) regions, defined as ±1 kb around the TSS, overlapped the interacting anchor, in each tissue. CRISPR-based evidence was used to link experimentally tested regulatory elements (like enhancers) to genes showing a significant perturbation response. Three tissue-agnostic combining strategies (cS2G, OpenTarget, and GeneHancer) (24-26) were incorporated using the source-provided V2G assignments, defined by confidence thresholds according to the original source. Together, these complementary data sources capture diverse mechanisms of gene regulation.

**Table 1.**
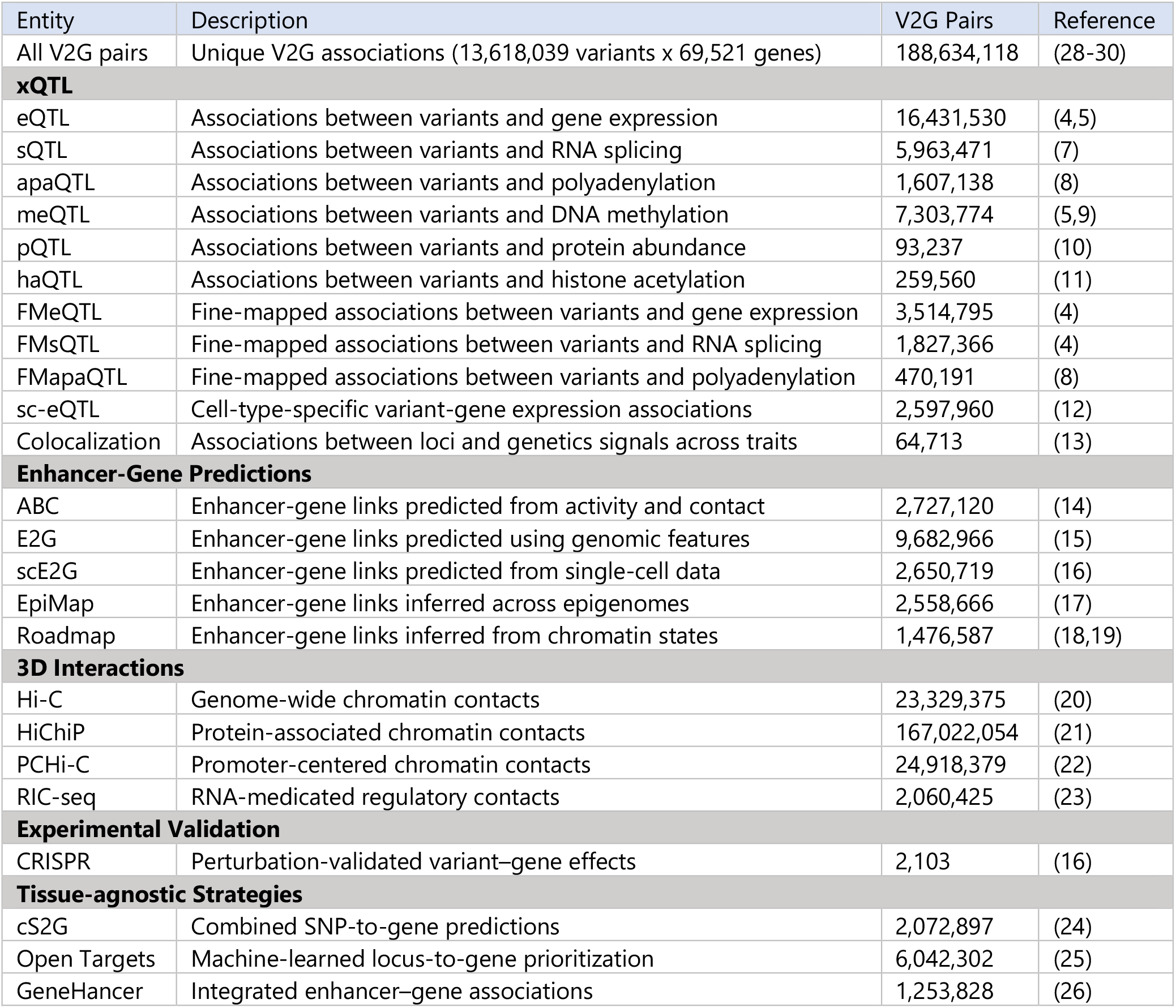
Summary of V2G pairs across all 24 evidence sources integrated into the mV2G atlas.

### V2G evidence integration for high-confidence V2G prioritization at tissue level

To enable data integration, all evidence datasets were harmonized to the V2G pair reference defined above and organized by V2G pair, tissue, and evidence type. For each V2G pair, the prioritization score, the number of evidence supporting its association, was counted independently for each tissue, allowing the same V2G association to receive different confidence levels across biological contexts. Tissue-specific evidence contributed only to its corresponding tissue, whereas tissue-agnostic evidence contributed across all tissues. After integration, 188,634,118 V2G pairs, involving 13,618,039 unique variants and 69,521 unique genes, are supported by at least one piece of evidence (prioritization score of at least 1) in one or more tissues.

Multiomic integration enables the identification of high-confidence functional V2G associations, as demonstrated in our previous study (27). To this end, we tested the prioritization score threshold using the fraction of prioritized variants with unique target genes. A prioritization score of at least 5 identifies only one target gene for more than 80% of prioritized variants in all tissue types, which we used to define high-confidence V2G associations. For each of these high-confidence V2G associations, we further defined their tissue specificity by whether they were shared across at least ten tissues. For high-confidence V2G pairs shared among multiple tissues, the top candidate tissue type was ranked based on the fraction of significant evidence relative to all available evidence in that tissue.

### Construction of mV2G atlas

The integrated data and precomputed tissue-specific scores were stored in an SQLite relational database containing standardized tables for variants, genes, tissues, evidence, and V2G pairs. Database indexing enabled rapid retrieval of information by variant, gene, tissue, or V2G pair for the mV2G atlas website. The web interface was implemented as a client-side JavaScript application, with a lightweight PHP backend exposing the SQLite database through a query API. User requests are sent to the server in the background, and results are displayed by updating the page directly without reloading it. Query results are rendered client-side as interactive tables supporting sorting, filtering, and pagination, and each entry links to a dedicated V2G annotation page. The embedded genome browser is aligned to the GRCh38/hg38 reference assembly and displays GENCODE v47 gene annotation, consistent with the GTEx v11 gene annotation used throughout the atlas (31,32). It integrates two functional-annotation tracks: the ENCODE SCREEN registry of candidate cis-regulatory elements (cCREs), which classifies elements as promoter-like, enhancer-like, or CTCF-bound signatures, and the Roadmap Epigenomics 15-state ChromHMM chromatin-state model, displayed for the epigenome matching the selected tissue (18,19,33,34).

## RESULTS

### Overview of the mV2G atlas

By integrating 21 tissue-specific evidence types and 3 tissue-agnostic prioritization strategies of V2G association types across 50 human tissues (**Fig. 1A**), the current release of the mV2G atlas has incorporated a total of 188,634,118 V2G pairs among 13,618,039 genetic variants and 69,521 genes, including 18,403 protein-coding genes and 51,118 noncoding RNAs, that are supported by at least one type of V2G association. The number of evidence-supported variants, genes, and V2G pairs varied across the 50 tissues, ranging from 8,952,481 in the brain amygdala to 133,370,096 in whole blood (**Fig. 1B**). This variation reflects differences in the coverage and functional interpretations of the underlying multiomic datasets, highlighting the importance of having tissue-specific results. By organizing all evidence within a tissue framework, mV2G allows users to compare the regulatory evidence landscape across tissues while preserving the biological context of each association.

**Figure 1.**
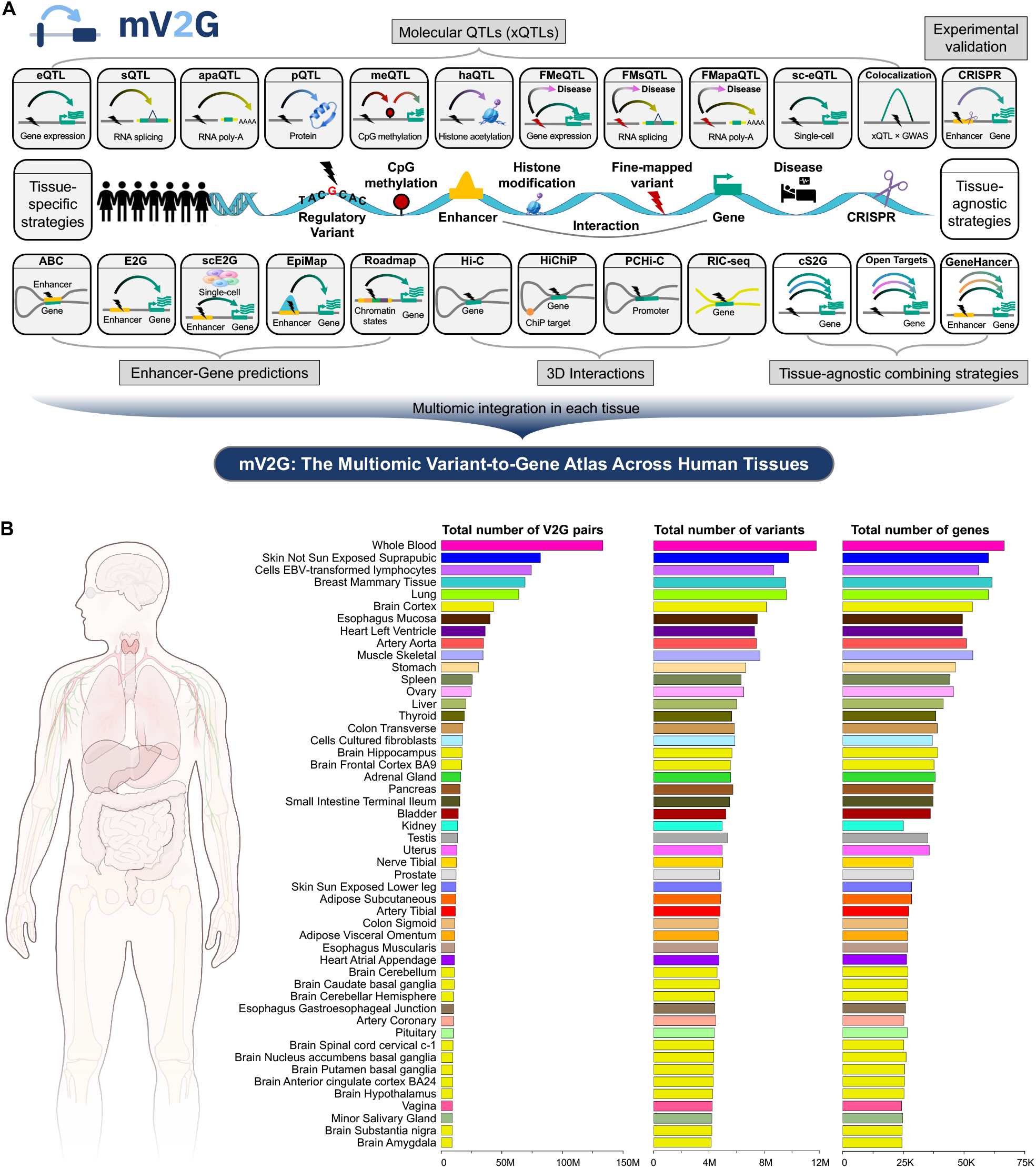
Overview of the mV2G atlas and its evidence landscape across tissues. (**A**) Schematic overview of the 24 V2G evidence types integrated into the mV2G atlas. The evidence sources include 21 tissue-specific evidence types grouped into xQTLs, enhancer– gene predictions, 3D chromatin interactions, and experimental validation, as well as 3 tissue-agnostic prioritization strategies. (**B**) Number of evidence-supported V2G pairs, unique variants, and unique genes in each tissue.

### Prioritization of high-confidence V2G associations

We evaluated the effect of increasing the prioritization score threshold on the number of retained V2G pairs as well as the number of variants targeting unique genes (**Fig. 2A**). As the prioritization score increased, the number of prioritized variants diminished. However, the proportion of variants targeting distinct genes showed a marked increase, indicating the effectiveness of our prioritization strategy in pinpointing authentic regulatory variants.

**Figure 2.**
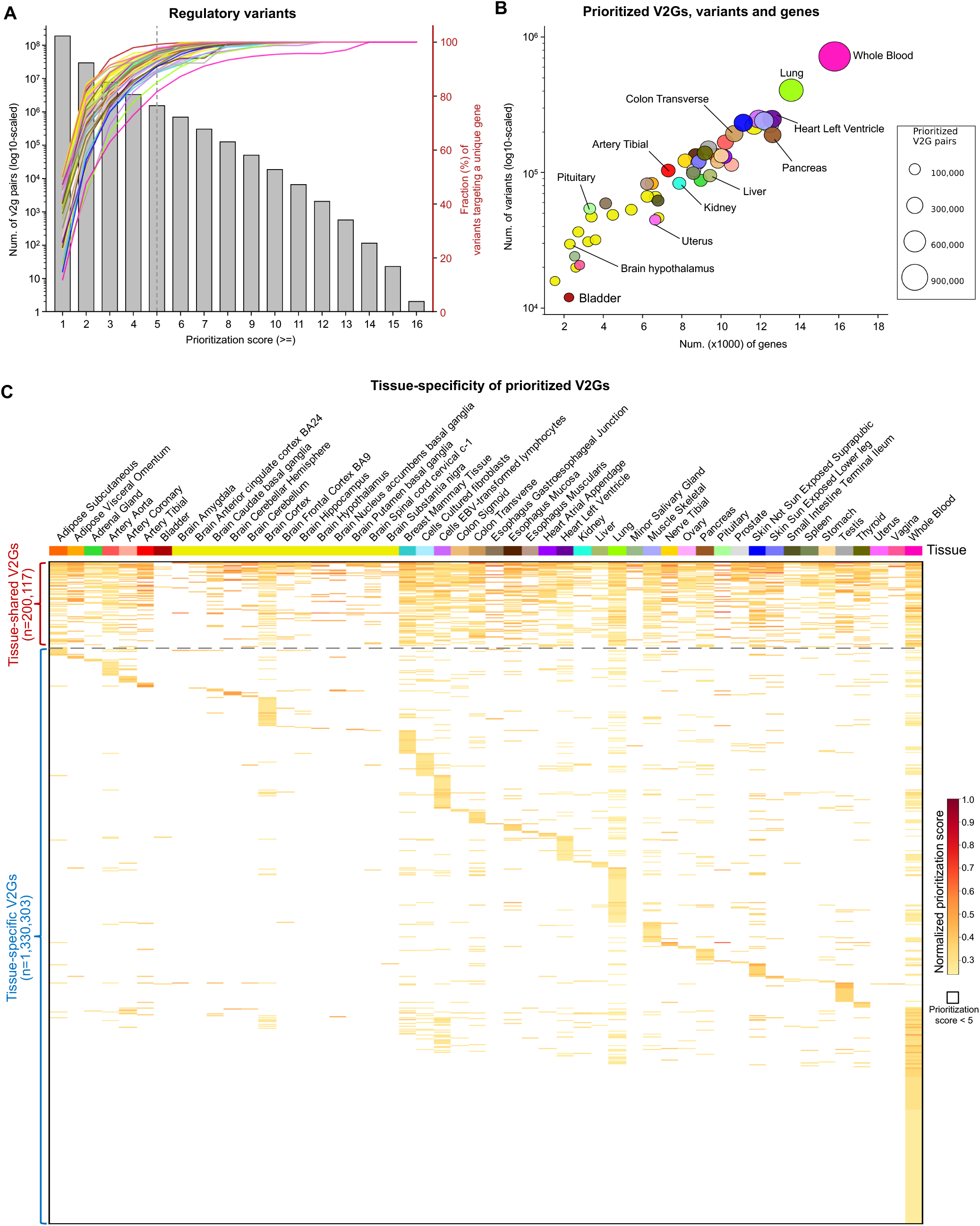
Prioritization and tissue specificity of high-confidence V2G associations. (**A**) Effect of increasing prioritization score thresholds on the V2G landscape. The bars show the number of V2G pairs retained at each score threshold on a log10 scale. The colored lines represent the 50 tissues and show the percent of variants linked to a uniquely prioritized candidate target gene at each threshold. The vertical dashed line indicates the selected high-confidence prioritization score threshold of ≥5. (**B**) Distribution of high-confidence V2G associations with a prioritization score of ≥5 in each tissue. Each point represents one tissue, and the point size represents the number of high-confidence V2G pairs. (**C**) Tissue-sharing and tissue-specificity patterns among high-confidence V2G associations.

Furthermore, it highlights the importance of large-scale evidence collection and integration to identify high-confidence V2G associations. Specifically, at score cutoff of 5, all 50 tissues had more than 80% their variants only associated with one gene. Based on these distributions, we selected a prioritization score of at least 5 to define high-confidence functional V2G associations. Although the tissues showed broadly similar trends, the separation among curves indicates differences in the amount and composition of regulatory evidence available in each biological context.

Requiring a prioritization score of at least 5 in one or more tissues identified 1,530,420 high-confidence functional V2G pairs involving 1,131,316 unique variants and 18,426 unique genes, including 16,469 protein-coding genes. Across the 50 tissues, this corresponded to an average of 139,988 high-confidence V2G pairs per tissue, ranging from 12,101 in the bladder to 908,639 in whole blood (**Fig. 2B**). These results demonstrate that the breadth of the prioritized V2G associations across tissues even after selecting for high-confidence associations, highlighting the importance of tissue-specificity.

### Tissue-specific V2G associations

Next, we examined the extent to which high-confidence V2G associations were specific to particular tissues or shared across multiple biological contexts (**Fig. 2C**). Most (87%) of high-confidence V2G associations were tissue-specific, while only 13% were shared across at least 10 tissues. These two aspects of tissue-specificity were both well captured in our database. For example, the variant rs7537004 was prioritized with high confidence for *CELA3B* in the pancreas alone, reaching a prioritization score of 10 with no high-confidence support in any other tissue (**Supplementary Fig. 1**). This strong, tissue-restricted signal is consistent with the biology of *CELA3B*, which encodes a chymotrypsin-like elastase whose expression is limited to the pancreas and whose variation has been linked to chronic pancreatitis and pancreatic cancer (35,36). In contrast, the variant rs2247303 was prioritized with high confidence for *RPL12* across all 50 tissues examined, reaching a score of at least 10 in 28 of them (**Supplementary Fig. 2**). This ubiquitous pattern of support reflects the housekeeping role of *RPL12*, which encodes a component of the 60S ribosomal subunit that is constitutively expressed across tissues (37,38). Together, these examples demonstrate that our prioritization framework captures both strictly tissue-specific and broadly shared regulatory relationships, and that the tissue in which a variant is prioritized carries genuine biological meaning. The tissue-sharing associations also revealed greater similarity among subsets of biologically related tissues like in the brain or blood-related tissues, while many other V2G associations showed more restricted patterns of support. These results emphasize that regulatory evidence for a V2G relationship cannot always be generalized across tissues and highlight the importance of retaining tissue context during V2G prioritization.

### Interactive exploration of V2G associations on the mV2G website

The mV2G atlas provides user-friendly and interactive interfaces for exploring genetic variants, genes, and V2G associations across 50 human tissues (**Fig. 3A**). Through the search engine, users can query variants of interest by rsID or GRCh38/hg38 genomic coordinates and genes of interest by gene symbol or Ensembl identifier. Variant-centered searches result in candidate target genes associated with the queried variant, whereas gene-centered searches display variants linked to the queried gene (**Fig. 3B, C**). Notably, we integrated an interactive genome browser to enable users to explore the V2G associations of interest within their local genomic context alongside genetic variants, functional genes, cis-regulatory elements, and tissue-specific chromatin states from resources including ENCODE and Roadmap (18,19,33,34). The interactive result tables summarize the candidate V2G associations, total evidence support, and highest-scoring tissue, allowing users to select a V2G association directly from a broad query to a dedicated V2G annotation page.

**Figure 3.**
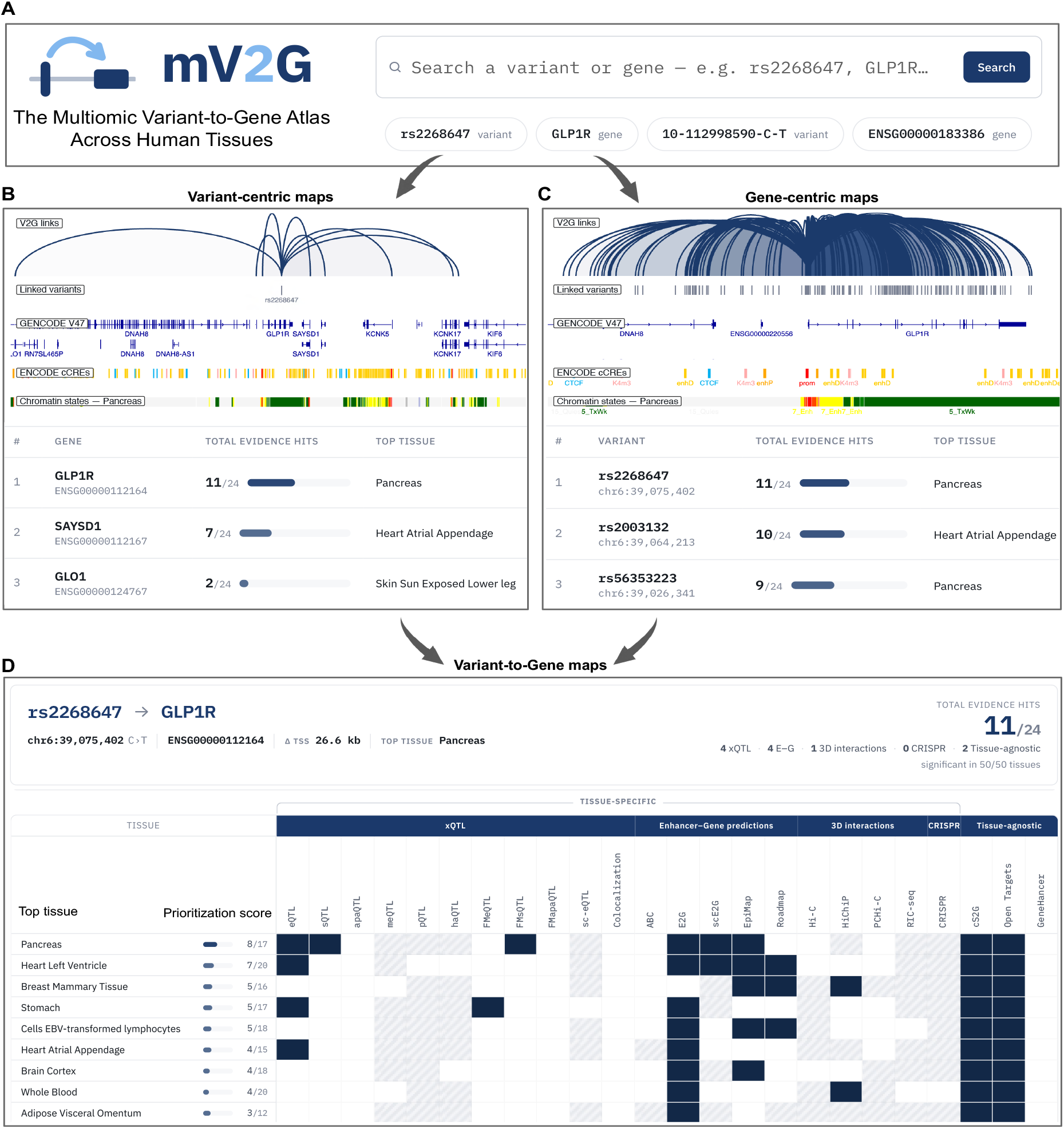
Example application of the mV2G atlas to prioritize and characterize the rs2268647-*GLP1R* association. (**A**) The mV2G atlas search interface enables queries by variant rsID, location, or a gene identifier. (**B**) Variant-centric view for rs2268647, displaying an IGV browser and a table to show top candidate target genes ranked by their integrated V2G evidence and the tissue with the strongest support for each association. (**C**) Gene-centric view for *GLP1R*, displaying an IGV browser and a table to show top regulatory variants linked to *GLP1R* and their corresponding evidence and top-supported tissues. (**D**) Detailed V2G view of the rs2268647-*GLP1R* association, showing prioritization scores across tissues and the individual evidence types supporting the association in each tissue.

For each V2G association, a dedicated annotation page provides a detailed tissue-resolved representation of the diverse evidence supporting this association (**Fig. 3D**). For each tissue, the interface displays the prioritization score together with the number of evidence types available in that tissue. An accompanying evidence matrix indicates support from individual xQTLs, enhancer–gene predictions, 3D chromatin interactions, experimental validation, and tissue-agnostic combining strategies. This design enables users to explore the tissue specificity of V2G associations and identify the top candidate tissue(s) in which a genetic variant affects the gene function, while retaining access to the individual evidence types underlying its prioritization. The resulting tissue-specific associations, evidence composition, and prioritization scores, can be downloaded in tabular formats for downstream analysis, data integration, and hypothesis generation.

### Case study: mV2G prioritizes rs2268647-*GLP1R* association in pancreas and heart

The rs2268647-*GLP1R* association illustrates how mV2G can prioritize a functional noncoding variant to a biologically and therapeutically relevant target gene in a tissue-specific context (**Fig. 3C**). *GLP1R* encodes the glucagon-like peptide-1 receptor, a major regulator of glucose-dependent insulin secretion, appetite, gastric emptying, and energy balance (39-41). GLP-1 receptor agonists have been used to treat type 2 diabetes and obesity (42). mV2G prioritizes *GLP1R* as the top candidate target gene of rs2268647, a variant located in the *GLP1R* intron and associated with body mass index (43). Notably, the rs2268647-*GLP1R* association receives its strongest support in the pancreas, with additional support in cardiac tissues. The pancreatic prioritization is supported by multiple complementary evidence classes rather than by a single dataset, providing an interpretable basis for the association. The strong support of the rs2268647-*GLP1R* association observed in the pancreas is consistent with the established role of *GLP1R* in glucose-dependent insulin secretion (44-46), while evidence in cardiac tissues is consistent with previous genetic and expression studies implicating *GLP1R* in cardiovascular biology (47-49). These findings provide genetic support for the clinical benefits of GLP-1 receptor agonists in patients with diabetes and cardiovascular disease (50). Together, this example demonstrates the utility of mV2G to prioritize tissue-specific variant-to-gene relationships supported by diverse regulatory evidence, facilitating the functional interpretation of noncoding variants and accelerating therapeutic target discovery.

## DISCUSSION

The mV2G Atlas addresses a central challenge in the interpretation of noncoding genetic variants by integrating diverse functional multiomic evidence to prioritize candidate target genes in a tissue-specific context. A key strength of our resource is its ability to preserve tissue information while combining complementary evidence derived from molecular QTLs, enhancer–gene predictions, 3D chromatin interactions, and experimental validation. Rather than assigning a single relationship to each variant and gene, the mV2G atlas allows users to evaluate how strongly an association is supported across different tissues and to examine the individual evidence underlying each prioritization. This framework may facilitate the interpretation of GWAS loci and the generation of testable hypotheses regarding the genes and tissues through which disease-associated noncoding variants exert their effects.

Preserving tissue specificity is a key feature of the mV2G Atlas. Because the regulatory effects of noncoding variants are highly tissue-dependent, a single variant may regulate distinct target genes across tissues or influence disease only within specific cellular contexts (3-5).

For example, mV2G prioritizes the rs2268647–*GLP1R* association in the pancreas and cardiac tissues, consistent with the established roles of GLP1R in glucose homeostasis, cardiovascular biology, and its clinical success as a therapeutic target (50). Therefore, assigning a single gene to a noncoding variant without considering tissue context may overlook biologically relevant regulatory mechanisms. By integrating diverse functional genomic evidence while preserving tissue context, the mV2G atlas facilitates tissue-specific V2G prioritization for disease mechanism interpretation and therapeutic target discovery.

As functional genomic resources continue to expand, future versions of the mV2G Atlas will incorporate additional datasets to increase the number and confidence of supported variant-to-gene associations. An important future direction will be to extend the current tissue-level framework to the resolution of individual cell types and cell states through the integration of single-cell and spatial genomic data (16,27,51). Incorporating additional disease-specific regulatory evidence and experimentally validated V2G relationships may further improve the identification of context-specific regulatory associations. Overall, by providing an accessible framework for integrating, prioritizing, and exploring V2G evidence across human tissues, the mV2G Atlas serves as an evolving resource for investigating the functional consequences of genetic variation and prioritizing candidate genes for downstream biological and experimental studies.

## ACKNOWLEDGEMENTS

We acknowledge the data support from the Functional Annotation of Variants Online Resources (FAVOR), the Impact of Genomic Variation on Function Consortium (IGVF), the GENCODE Project, the Genotype-Tissue Expression (GTEx), the Encyclopedia of DNA Elements (ENCODE), the Roadmap Epigenomics, and other studies. We thank the Center for Integrated Research Computing (CIRC) at the University of Rochester for computational support.

## AUTHOR CONTRIBUTIONS

Conceptualization: H.L. and D.Z.; Data curation, Formal analysis, Investigation, Methodology, Project administration, Software, Validation, Visualization: D.Z.; Data resources: H.Z., X.L., M.U.S., A.R.G., and J.M.E; Project administration and Supervision: H.L.; Writing– original draft: D.Z. and H.L. Writing—review & editing: D.Z., H.L., H.Z., M.U.S., A.R.G., J.M.E, and X.L.

## SUPPLEMENTARY DATA

Supplementary Data are available at NAR online.

## DATA AVAILABILITY

The mV2G Atlas and the processed data described in this article are freely available at https://mv2g.hbliulab.org/. Data can be queried and downloaded through the database website. All third-party datasets integrated into the mV2G Atlas were obtained from publicly available resources, as described and cited in the article, and remain available through their respective source repositories.

