## Supplementary figures and images for "mV2G: a multiomic atlas for tissue-specific variant-to-gene prioritization"

### Supplementary Figures

#### Supplementary Figure 1

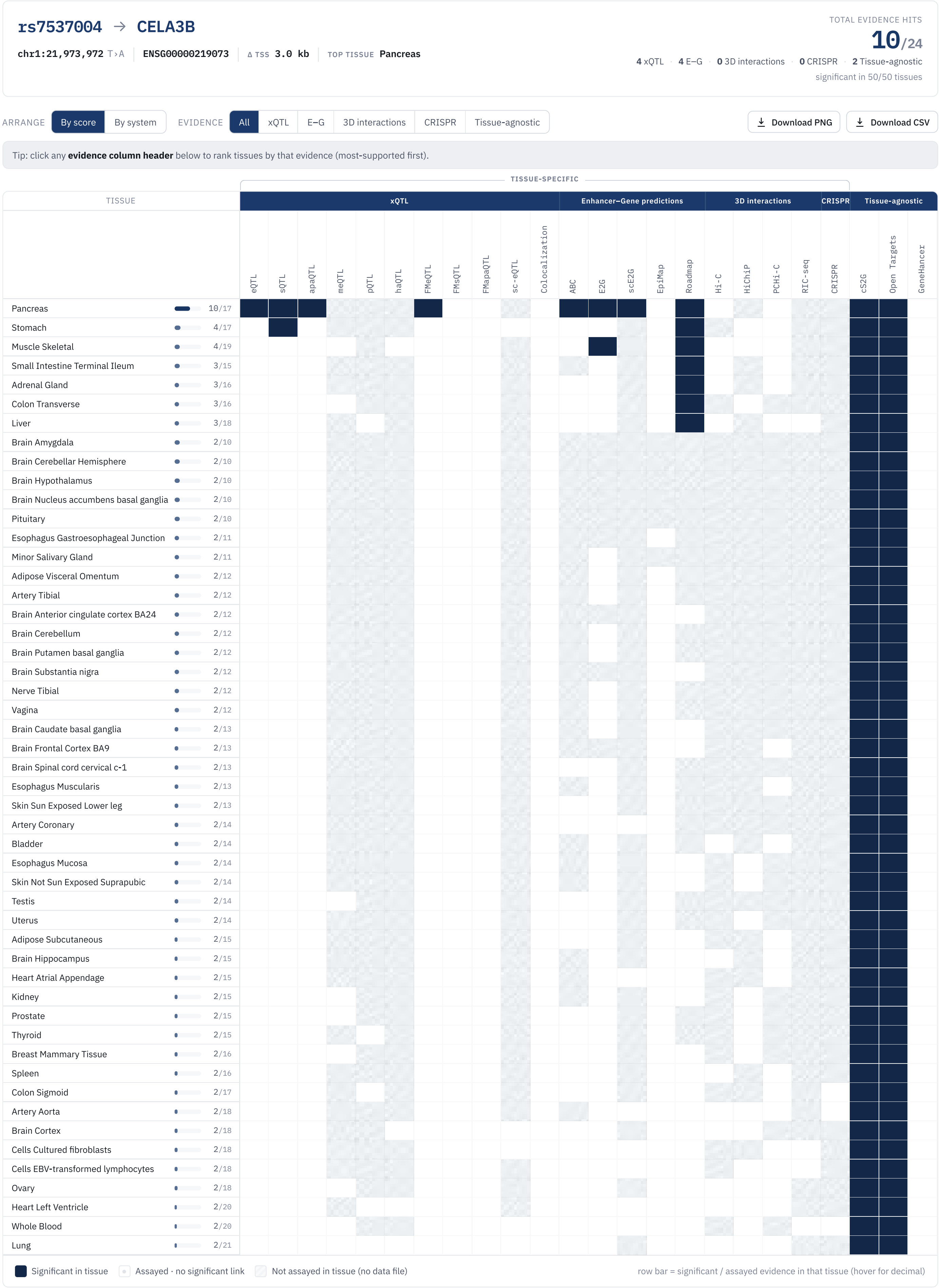

#### Supplementary Figure 2

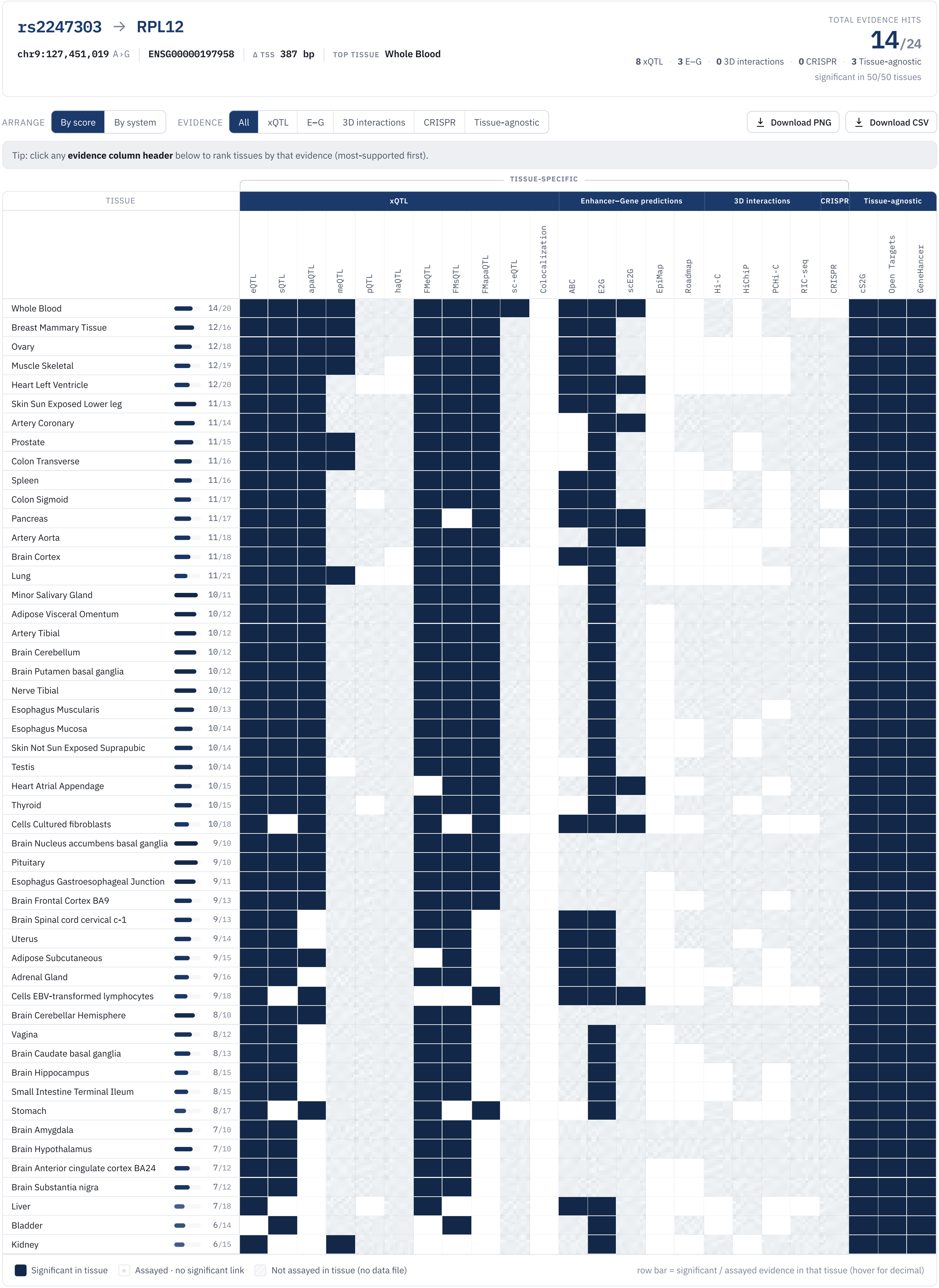
